# Genome-wide association study of DMI fungicide sensitivity detects numerous small-effect variants in the major North American population of *Fusarium graminearum*

**DOI:** 10.1101/2025.09.25.678624

**Authors:** Upasana Dhakal, Christopher Toomajian

## Abstract

Fusarium head blight (FHB), a major disease of wheat, is primarily managed through applications of demethylation inhibitor (DMI) fungicides during anthesis. However, repeated use of DMIs has led to the emergence of *Fusarium graminearum* isolates with reduced sensitivity and, in some cases, resistance. In this study, we evaluated the sensitivity of 152 *F. graminearum* isolates to propiconazole and tebuconazole. While sensitivity varied among isolates, no resistant strains were detected. We also conducted a genome-wide association study (GWAS) to investigate the genetic basis of DMI sensitivity. GWAS identified 48 and 39 quantitative trait nucleotides (QTNs) associated with propiconazole and tebuconazole sensitivity, respectively, with 12 QTNs common to both fungicides—supporting their common mode of action. Candidate gene analysis highlighted genes encoding transporters, secondary metabolite synthesis enzymes, transcription factors, and a heat shock protein as potential candidates for DMI fungicide response. We propose that tolerance to DMIs in *F. graminearum* is linked to active fungicide efflux out of fungal cells, mediated by transporters, including those associated with secondary metabolite pathways.

**Data summary:** Information on all strains used in the experiments have been included in a supplemental file. All sequencing data used in this study are publicly available and were described in our previous publication, which has been cited. Additional supporting data (isolate genotype and phenotype files) and code written for the analyses described here are made available on GitHub: https://github.com/Toomajian-laboratory/files_fungicide_manuscript/

**Impact statement:** Isolates from field populations of fungal plant pathogens vary in their sensitivity to DMI fungicides, though in most cases the genetic determinants of this variation are poorly understood. In this study, we measure the sensitivity of isolates from the main US population of *Fusarium graminearum* to two DMI fungicides. We used GWAS to identify genomic loci and candidate genes that might underlie variation in DMI sensitivity. Our work contributes to the broader effort to understand the evolution of fungicide tolerance and resistance in populations of plant pathogens. The candidate genes identified, most of which are novel, provide good targets for functional studies of the tools fungi use to survive fungicides. The identified variants can also be screened to monitor potential increases in fungicide tolerance. The high-throughput, rapid, and large-scale monitoring of fungicide sensitivity described here advances both fundamental research and resistance management efforts.

## Introduction

FHB is one of the major diseases of wheat in North America and other wheat-growing areas of the world. In North America, *F. graminearum* is the major causal agent of FHB (Goswami and Kistler, 2004). Wheat heads are susceptible to infection from anthesis to the soft dough stage and are most susceptible at anthesis. Infected heads become prematurely bleached. Grains from infected heads are shriveled, discolored, and lightweight. When the weather conditions are favorable (high humidity during anthesis), FHB can destroy the crop in a matter of a few weeks (McMullen et al., 1997).

In addition to cultural management practices like growing resistant cultivars, crop rotation, and crop residue management, fungicide application during anthesis is an important component of integrated FHB management. DMI fungicides like propiconazole, metconazole, prothioconazole, and tebuconazole have been used to manage FHB in the US since the mid-1990s (McMullen et al., 2012). DMI fungicides inhibit the synthesis of ergosterol, an important component of fungal cell walls. Members of the CYP51 gene family in fungi are required for ergosterol biosynthesis. DMI fungicides inhibit ergosterol synthesis by binding to 14-a demethylases, the product of the CYP51 genes. *Fusarium graminearum* has three CYP51 paralogs, *CYP51A*, *CYP51B*, and *CYP51C*, encoding 14-a demethylases (Liu et al., 2011).

Intensive application of these fungicides has led to the development of fungicide resistance in some *F. graminearum* field isolates/populations collected from Asia, Europe, and the USA (Klix et al., 2007; Spolti et al., 2014; Yin et al., 2009). Application of a mixture of two or more DMI fungicides is a common practice in disease management. Cross-resistance among DMI fungicides has also been reported in several studies. For example, isolates exhibiting cross-resistance between tebuconazole and metconazole were documented by (Spolti et al., 2012), and additional cross-resistance among metconazole, tebuconazole, ipconazole, epoxiconazole, and difenoconazole was observed by (Duan et al., 2018). In contrast, (Liu et al., 2022) found no evidence of cross-resistance between prothioconazole and tebuconazole. Resistance to azole fungicides, a specific class within the broader DMI fungicide category, has been reported in other fungal species besides *F. graminearum*. Resistance to azoles has occurred through multiple mechanisms, such as mutation in CYP51 genes, overexpression of CYP51 genes, and transport of fungicide out of fungal cells. Mutations in CYP51 genes are associated with azole fungicide resistance in the fungus *Mycosphaerella graminicola* (Leroux et al., 2007). Repeated sequences upstream of CYP51 genes resulting in overexpression of CYP51 genes have been identified in resistant isolates of *Blumeriella jaapii* and *Penicillium digitatum* (Hamamoto et al., 2000; Ma et al., 2006). In addition to overexpression, transport of fungicides out of the fungal plasma membrane through efflux pumps is another mechanism conferring fungicide resistance in fungi (Hayashi et al., 2003; Reimann and Deising, 2005). Different DMI fungicides have differential binding preferences to different CYP51 paralogs in *F. graminearum* (Liu et al., 2011). *CYP51A* disruption mutants were sensitive to seven DMIs including propiconazole and triadimefon whereas *CYP51C* disruption mutants were sensitive to tebuconazole and four other DMIs but not to propiconazole (Liu et al., 2011).

In field isolates, non-synonymous mutations in *CYP51A* and *CYP51C* genes were not correlated to DMI sensitivity, likely due to the contribution of genes outside the CYP51 gene family to DMI tolerance (Talas and McDonald, 2015). Thay also did not find any mutations likely regulating the overexpression of genes in the promoter regions for all three *F. graminearum* CYP51 paralogs. They did, however, find high variability and heritability (0.97) in fungicide sensitivity among *F. graminearum* isolates. This opens the possibility of using genome-wide association (GWAS) to identify the genetic basis of variation in fungicide sensitivity in *F. graminearum*. This approach is free of any hypothesis and screens the whole genome without any bias to a specific region or type of gene or variant (Kitsios and Zintzaras, 2009), allowing for the identification of previously unknown genes and pathways related to fungicide sensitivity (Manolio et al., 2008).

Two studies have applied GWAS in *F. graminearum* populations to identify the contribution of genes outside the CYP51 gene family to DMI sensitivity. (Talas et al., 2016) performed GWAS for propiconazole sensitivity with a sample of 220 strains collected in 2008 from German field populations that had been genotyped with a reduced representation shotgun sequencing approach. They report 74 SNPs significant at a false discovery rate (FDR) of 0.05, with three of these significant using a more conservative Bonferroni threshold. (Poudel et al., 2024) performed GWAS for sensitivity to both tebuconazole and prothioconazole in a sample of 183 isolates from North Dakota collected over several decades, from 1981 to 2013, and identified 7 significant marker-trait associations.

Here, we use GWAS to identify genetic determinants of sensitivity to both propiconazole and tebuconazole in field isolates of *F. graminearum*. We use isolates that have been genotyped by genotyping by sequencing (GBS) in a broader population genomic study of multiple populations from the Americas (Dhakal et al., 2024). However, for the GWAS we selected isolates almost exclusively from the North America 1 (NA1) population that were collected from 1999 to 2013 from multiple regions of the US. Our study aims to extend prior GWAS of DMI sensitivity in *F. graminearum* by using a different sample that focuses on the major population found in North America.

## Methods

### Fungal isolates

One hundred and fifty-two *Fusarium graminearum* isolates were used for this study. The inference of *TRI* loci genotype and population membership for each isolate were described previously (Dhakal et al., 2024). These strains were collected from Minnesota, Montana, North Dakota, New York, Louisiana, and Virginia by other researcher (Burlakoti et al., 2011; Gale et al., 2011, 2007; Kuhnem et al., 2015; Puri and Zhong, 2010; Spolti et al., 2014; Zeller et al., 2004). Isolates were collected between 1999 and 2013, with most of them collected in the years 2000 (76) and 2013 (43) (Supplemental Table 1).

### Genotyping

Isolates used here are a subset of the isolates used in our previous population genomics study (Dhakal et al., 2024). Isolates were sequenced using genotyping by sequencing (GBS) (Yue, 2017). Polymorphisms were extracted using GATK 4.1.8.1 (McKenna et al., 2010). Due to relatively high levels of missing genotype call data from the GBS process, we performed imputation of missing genotype calls using most of the isolates from our larger study. We started with genotype calls from 555 isolates, the original 570 isolates sequenced in that study minus the six with the fewest genotype calls and an additional nine that differed from the PH-1 isolate at over 10% of the identified SNPs. SNPs with genotype calls in less than 25% of the set of 555 isolates were discarded, and the remaining 63,102 SNPs were used in the imputation process. Missing sites were imputed for the set of 555 isolates using Beagle 5.1 (Browning and Browning, 2007). After imputation, the SNPs were filtered to exclude sites with minor allele frequency less than 0.05 in the set of 152 isolates used in the present study, leaving 4,857 SNPs for subsequent analysis.

### Fungicide sensitivity assay

Sensitivity of 152 *F. graminearum* isolates towards two DMI fungicides, propiconazole and tebuconazole, was tested using a flat bottom 96 well plate assay (Talas and McDonald, 2015). Propiconazole and tebuconazole were tested in 8 different concentrations per experiment. For propiconazole, the following concentrations were used for all three replications: 701 *μ*M, 234 *μ*M, 87.7 *μ*M, 29.2 *μ*M, 13.1 *μ*M, 4.38 *μ*M, 1.46 *μ*M, and 0 *μ*M. For the first replication for tebuconazole, the following concentrations were used: 78.0 *μ*M, 26.0 *μ*M, 9.75 *μ*M, 3.25 *μ*M, 1.46 *μ*M, 0.487 *μ*M, 0.162 *μ*M, and 0 *μ*M while the second and third replication used the following concentrations: 702 *μ*M, 234 *μ*M, 78.0 *μ*M, 26.0 *μ*M, 9.75 *μ*M, 3.25 *μ*M, 1.46 *μ*M, and 0 *μ*M. Fungicide solutions were prepared by mixing the fungicide with synthetic nutrient broth (Talas and McDonald, 2015). The concentration of DMSO was kept constant at 1.95 % (V/V) for all fungicide concentrations. Streptomycin sulphate was added to the liquid media at the rate of 0.33 mg/ml to prevent bacterial contamination. 150 µl of fungicide solution and 50 µl spore suspension (adjusted to 1×10^5^ spores/ml) were added to the wells of PR1MA flat bottom 96 well plates (MIDSCI) using a liquid handling robot (Beckman Coulter, Biomek NX^p^). All of the reported concentrations (fungicide, DMSO, and streptomycin) are for the final solution (150 µl fungicide solution + 50 µl spore suspension). Control wells had water rather than a spore suspension added to the fungicide and media solution. Spores were collected by growing *F. graminearum* isolates in 40 ml mung bean broth for about 8 days in 100 ml conical flasks. Mycelia was filtered out with double layered muslin cloth, and the liquid culture was centrifuged at 4000 rpm for 5 minutes to get a pellet of macroconidia in 50 ml centrifuge tubes. Liquid media was carefully removed without disturbing the pellet and spores were resuspended in sterile double distilled water, counted using a hemocytometer and frozen at -80°C. Tubes with concentrated spore solutions were thawed overnight and adjusted to the required concentration on the day of the inoculation. The plates were covered with lids, sealed with parafilm and incubated in the dark at room temperature for seven days. To provide humid conditions, plates were incubated over a mat of moist paper towels or incubated near water-filled trays. Absorbance readings were taken immediately after preparing the plates, and at 5 and 7 days at 405 nm using a plate reader (FLUOstar Omega SNP, BMG LABTECH). Isolates were evaluated using a complete block design, and the experiment was repeated twice, resulting in a total of three replicates.

### Half-maximal effective concentration (EC_50_) estimation

Separately for each fungicide × isolate × replicate combination, our dose-response data of absorbance readings of plate cultures corresponding to each fungicide concentration were fitted to a four parameter logistic model using R package dr4pl (An et al., 2019). The four estimated parameters included effective concentration 50 (EC_50_), upper limit, lower limit, and slope. For the first replication of both fungicide experiments, one of the 152 isolates was not included in the experiments. Additionally, for the first replication of the tebuconazole experiments, the EC_50_ values of 17 isolates were dropped when the estimated parameters suggested a poor model fit, specifically, when the lower limit parameter of the dose-response curve was more negative than -0.01. Similarly, for the first replication of the propiconazole experiment, the EC_50_ values for 24 isolates included on 2 of the experimental plates were excluded since for 23 of these 24 isolates, the lower limit parameter estimate was unusually high, greater than 0.22. Least squares (LS) means were calculated across the three replicates using a complete block design where isolate effect was modeled as a fixed effect and replicate as a random effect.

### Estimation of broad sense heritability

Broad-sense heritability was estimated using variance components of a linear mixed model in PROC MIXED (SAS Institute), treating isolates and blocks as random effects. Heritability was calculated as the ratio of genetic variance (variance related to isolate genotype) to the total phenotypic variance (variance related to isolate genotype, replicates, and residual variances).

### GWAS analysis

Genome wide association analyses were conducted using multi-locus mixed linear models implemented in R package `mrMLM` 5.0.1 (Zhang et al., 2020). Five mrMLM 5.0.1 models, namely mrMLM, FASTmrMLM (FAST mrMLM multi-locus mixed linear model), FASTmrEMMA (fast multi-locus random-SNP-effect EMMA), pKWmEB (integrates Kruskal-Wallis test with empirical Bayes), and ISIS EM BLASSO (iterative modified-Sure Independence Screening EM-Bayesian Lasso) were used. The models from the `mrMLM` R package perform multi-locus genome-wide association analysis where the effects of all markers are estimated simultaneously. They work in two steps. In the first step, each SNP is tested individually for association, treating marker effects as random, and a relaxed Bonferroni threshold is applied to select potentially associated markers. In the second step, a multi-locus mixed linear model estimates the effects of these selected markers using an empirical Bayes approach implemented via the Expectation-Maximization (EM) algorithm. True quantitative trait nucleotides (QTN) were confirmed with a likelihood ratio test, with LOD scores greater than or equal to 3 considered significant. No population structure was included in the models.

### Candidate gene analysis

As the genotypes used in the GWAS analysis come from GBS data that represent a fraction of the total genome, associated SNPs are often likely markers in linkage disequilibrium (LD) with nearby causative variants that were not captured directly by GBS. Thus, for each associated SNP, we define a QTN region within which candidate genes are highlighted. Our previous work indicates that within the NA1 population, LD decays on average within about 3 kb. As a default, each QTN region was defined as the window 3 kb in either direction of the associated SNP (or 6,001 bp in total). Additionally, for each QTN region, its boundaries were occasionally adjusted after manually inspecting the LD decay pattern on both sides of the QTN. The squared correlation coefficient (*r*^2^) pairwise LD measure between the QTN and SNPs close to it on the left and right were used to adjust left and right QTN region boundaries. For five QTN regions, the default boundaries 3 kb away from the associated SNP were expanded due to the presence of a nearby SNP in high LD with the associated SNP, where pairwise *r*^2^ >0.75. The regions were expanded in units of kb to approximate the distance to any linked SNPs in high LD. Total region size is the sum of left and right LD blocks. Genes falling in the QTN regions were obtained using a custom python script. The curated phenotypes and gene ontology (GO) terms for the candidate genes were obtained from FungiDB (https://fungidb.org/fungidb/app).

## Results

### Sample composition, population assignment and linkage disequilibrium decay

As our main goal was to investigate variation in fungicide sensitivity in the North American 1 (NA1) population of *F. graminearum*, we chose the 152 isolates, primarily from the NA1 population, for inclusion in this study from a larger set of diverse *F. graminearum* isolates characterized in our previous population genomics study (Dhakal et al., 2024). This previous study describes variant calling from GBS reads and the determination of *TRI* loci genotypes and population membership for the isolates used in this study (Dhakal et al., 2024). Of the 152 isolates analyzed, 151 had the 15ADON *TRI* genotype, and 147 were assigned to the NA1 population (Dhakal et al., 2024). Most of the isolates were collected from three states: New York (73), Minnesota (35), and North Dakota (33), while a much smaller proportion of samples came from Montana (8), Louisiana (1), Michigan (1) and Virginia (1). Sampling occurred across several years: 1999 (9 isolates from MN and ND), 2000 (61 isolates from MN, MT, ND, and NY), 2003 (1 isolate from LA), 2008 (3 isolates from ND and NY), 2010 (1 isolate from VA), 2011 (16 isolates from NY), and 2013 (43 isolates from NY). In the larger sample of the NA1 population that we have investigated, linkage disequilibrium (LD) decays to half of its maximum within 3 kb, suggesting that GWAS in this population could achieve high-resolution mapping (Dhakal et al., 2024).

### Fungicide sensitivity

Isolates displayed variation in their sensitivity to propiconazole and tebuconazole. Yet none of our isolates were obviously resistant to propiconazole and tebuconazole, as the least sensitive isolate to each fungicide had an EC_50_ value only about three times as great as the sample average. The EC_50_ values for the two fungicides approximate a normal distribution with the addition of a long right tail (Figure 1). The EC_50_ values ranged from 8.3 *μ*M to 100 *μ*M for propiconazole and 4 *μ*M to 49 *μ*M for tebuconazole (Figure 1). Isolate 23310 collected in 2013 from Belmont, NY was the most tolerant to both fungicides, whereas isolate 24315 collected in the year 2000 from Logan County, North Dakota was most sensitive to both fungicides. Both of these isolates have the 15ADON *TRI* loci genotype. Tebuconazole was more effective in suppressing fungal growth, as the average EC_50_ was lower for tebuconazole (17 *μ*M) than for propiconazole (35 *μ*M). Propiconazole and tebuconazole sensitivities were highly positively correlated, with a Pearson correlation coefficient of 0.896. The broad sense heritability for propiconazole sensitivity and tebuconazole sensitivity are 0.84 and 0.8, respectively.

**Figure 1:**
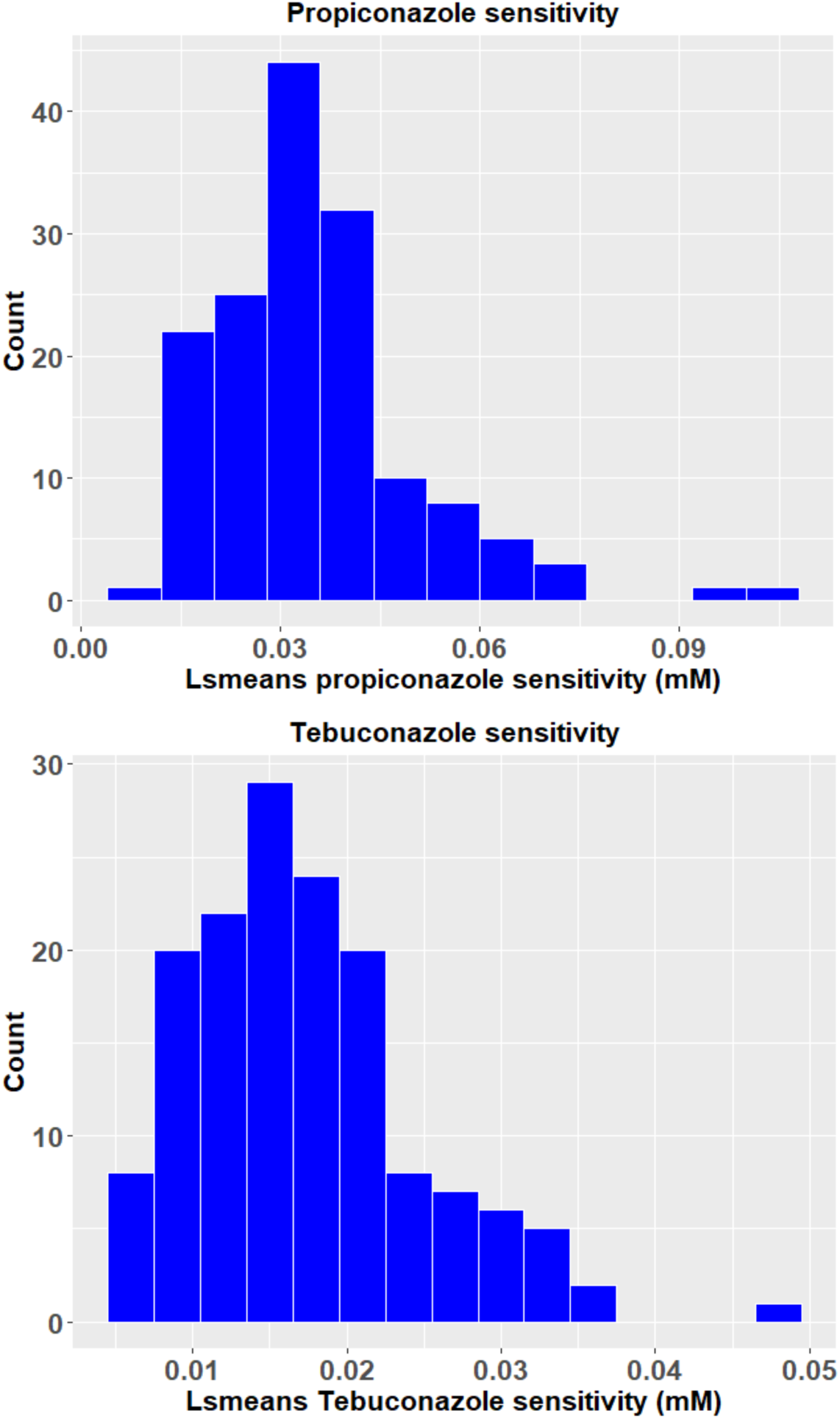
Distribution of least squares means of EC_50_ for propiconazole and tebuconazole.

### GWAS

GWAS analysis for the traits propiconazole and tebuconazole sensitivity was performed by using EC_50_ values estimated for 152 isolates and 4,857 SNPs distributed across the genome using models implemented in the software package mrMLM (Zhang et al., 2020). A kinship matrix was used to account for the relatedness among the individuals. Population structure adjustment was not necessary, as the vast majority of isolates in our sample came from the well-defined NA1 population (Dhakal et al., 2024), and a principal components analysis of these isolates does not identify clear subclusters (Supplemental Figure 1). Additionally, preliminary forward model selection in GAPIT version 3 using the Bayesian information criterion indicated that the optimal number of principal components to be included in its mixed linear model (MLM) was zero, consistent with no population structure. Diagnostic quantile-quantile (Q–Q) plots also verified that any trace population structure in the sample was not inflating P-values. Models in the mrMLM package identified 48 distinct significant QTNs for propiconazole sensitivity with LOD scores ≥ 3, 2 detected by the model mrMLM, 1 by FASTmrMLM, 30 by ISIS EM-BLASSO, and 22 by pKWmEB (Figure 2, Table 1). Six QTNs were detected by more than one method (Table 1). Even though many QTNs were detected, the majority of them (36) are each estimated to explain less than 1% of the total phenotypic variation in all models that identified them as significant (Table 1). QTN 6640556 on chromosome 3 explained the highest percent (10%) of the total phenotypic variation under the mrMLM model (Table 1). Similarly, models implemented in the package mrMLM detected 39 significant QTNs for the fungicide tebuconazole, 22 by the model ISIS EM-BLASSO and 17 by pKWmEB (Figure 3, Table 2). Two of these QTNs were detected by both models. Similar to propiconazole, 25 of the detected tebuconazole QTNs each explain less than 1% of the total variation in all models that identified them as significant. The highest phenotypic variation (3.06%) was explained by QTN 8322036 on chromosome 2. There are 13 QTNs common between propiconazole and tebuconazole (Tables 1 and 2, Supplemental Table 2).

**Figure 2:**
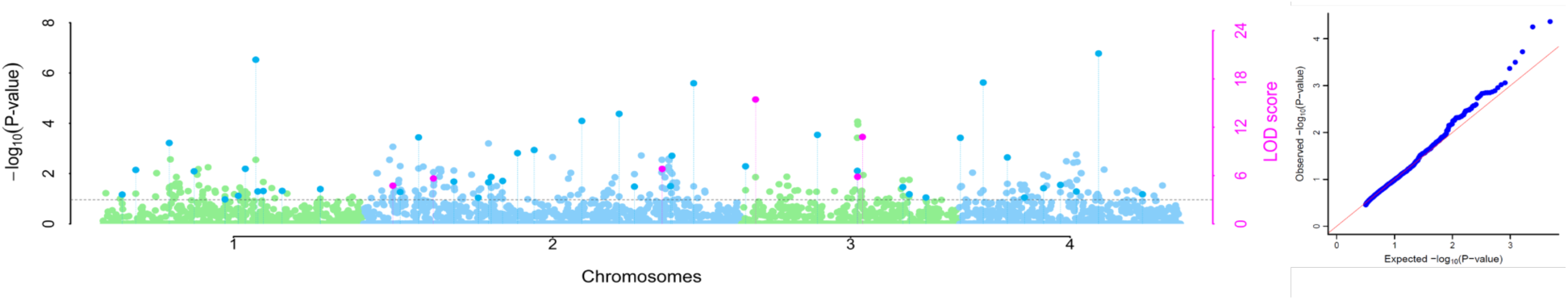
Manhattan and quantile-quantile plots for propiconazole sensitivity. Analysis was performed, and plots generated, using models mrMLM, FASTmrMLM, FASTmrEMMA, ISIS-EMBLASSO, and pKWmEB implemented in the mrMLM package. QTNs with significant LOD scores have dotted vertical lines that drop down to the x-axis. Those significant QTNs detected by a single model are shown in blue, while QTNs identified by two or more models are shown in pink. The remaining points represent the median of the intermediate -log₁₀(P-values) from mrMLM, FASTmrMLM, FASTmrEMMA, and pKWmEB.

**Figure 3:**
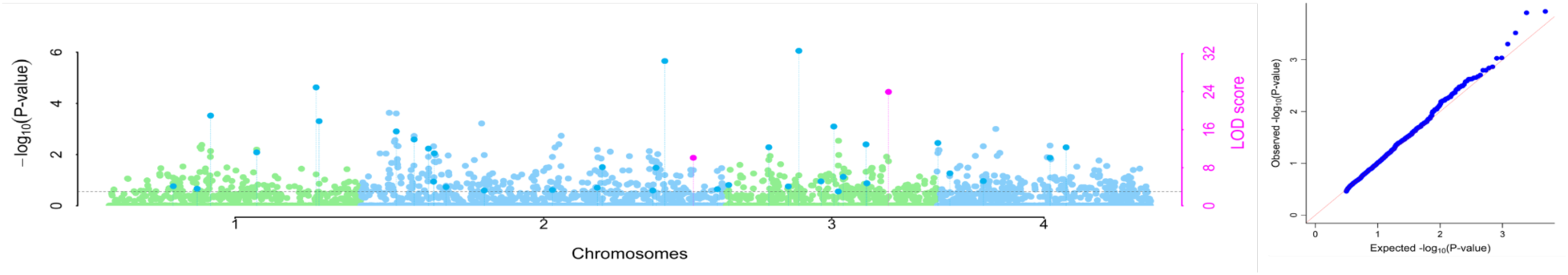
Manhattan and quantile-quantile plots for tebuconazole sensitivity. Analysis was performed, and plots generated, using models mrMLM, FASTmrMLM, FASTmrEMMA, ISIS-EMBLASSO, and pKWmEB implemented in the mrMLM package. QTNs with significant LOD scores have dotted vertical lines that drop down to the x-axis. Those significant QTNs detected by a single model are shown in blue, while QTNs identified by two or more models are shown in pink. The remaining points represent the median of the intermediate -log₁₀(P-values) from mrMLM, FASTmrMLM, FASTmrEMMA, and pKWmEB.

**Table 1.** List of QTNs significantly associated with propiconazole.

| Method | Chromosome | Marker position (bp) | QTN effect | LOD score | '-log10(P)' | r2 (%) | MAF |
| --- | --- | --- | --- | --- | --- | --- | --- |
| pKWmEB | 1 | 237393 | -2.00E-04 | 3.64 | 4.37 | 0.02 | 0.10 |
| ISIS EM-BLASSO | 1 | 489860 | -7.00E-04 | 6.70 | 7.55 | 0.11 | 0.13 |
| pKWmEB | 1 | 3881005 | 4.00E-04 | 10.05 | 10.98 | 1.49 | 0.34 |
| ISIS EM-BLASSO | 1 | 5731674 | -2.00E-04 | 6.52 | 7.37 | 0.01 | 0.09 |
| pKWmEB | 1 | 6829683 | -2.00E-04 | 3.04 | 3.73 | 0.06 | 0.23 |
| ISIS EM-BLASSO | 1 | 7218760 | -2.89E-05 | 3.48 | 4.20 | 0.00 | 0.38 |
| ISIS EM-BLASSO | 1 | 7356780 | -2.00E-04 | 6.83 | 7.69 | 0.00 | 0.06 |
| ISIS EM-BLASSO | 1 | 7467655 | -0.0075 | 20.37 | 21.46 | 5.26 | 0.05 |
| ISIS EM-BLASSO | 1 | 7514533 | -1.00E-04 | 4.02 | 4.78 | 0.01 | 0.29 |
| ISIS EM-BLASSO | 1 | 7585853 | 6.00E-04 | 4.08 | 4.83 | 0.12 | 0.28 |
| ISIS EM-BLASSO | 1 | 9999192 | 1.00E-04 | 4.09 | 4.85 | 0.01 | 0.31 |
| pKWmEB | 1 | 11270238 | 6.00E-04 | 4.31 | 5.08 | 0.01 | 0.22 |
| mrMLM | 2 | 459782 | -0.0041 | 3.58 | 4.30 | 8.11 | 0.48 |
| ISIS EM-BLASSO | 2 | 459782 | -0.0018 | 5.91 | 6.74 | 1.59 | 0.47 |
| pKWmEB | 2 | 501355 | -5.00E-04 | 3.95 | 4.70 | 0.28 | 0.10 |
| ISIS EM-BLASSO | 2 | 665406 | 0.0019 | 10.74 | 11.69 | 1.44 | 0.30 |
| FASTmrMLM | 2 | 851037 | 0.0031 | 3.32 | 4.04 | 3.27 | 0.24 |
| pKWmEB | 2 | 851037 | 1.60E-03 | 32.51 | 33.70 | 1.86 | 0.24 |
| ISIS EM-BLASSO | 2 | 851037 | 0.0027 | 5.64 | 6.46 | 2.48 | 0.24 |
| pKWmEB | 2 | 2753268 | 1.40E-03 | 5.25 | 6.05 | 0.54 | 0.11 |
| ISIS EM-BLASSO | 2 | 3804766 | -7.00E-04 | 3.24 | 3.95 | 0.20 | 0.50 |
| ISIS EM-BLASSO | 2 | 3923133 | -0.0025 | 5.16 | 5.96 | 0.83 | 0.08 |
| ISIS EM-BLASSO | 2 | 3938384 | 0.0014 | 5.82 | 6.64 | 0.79 | 0.33 |
| pKWmEB | 2 | 4138058 | -4.00E-04 | 5.33 | 6.13 | 0.89 | 0.13 |
| ISIS EM-BLASSO | 2 | 4358147 | 1.00E-04 | 8.79 | 9.70 | 0.00 | 0.05 |
| ISIS EM-BLASSO | 2 | 4679670 | -7.83E-05 | 9.16 | 10.08 | 0.00 | 0.06 |
| ISIS EM-BLASSO | 2 | 5664783 | -0.0013 | 12.77 | 13.76 | 0.80 | 0.44 |
| pKWmEB | 2 | 6243290 | -8.06E-05 | 13.66 | 14.66 | 1.01 | 0.31 |
| pKWmEB | 2 | 6487389 | 2.00E-04 | 4.62 | 5.40 | 0.40 | 0.13 |
| pKWmEB | 2 | 8062916 | -2.00E-04 | 8.86 | 9.77 | 0.01 | 0.39 |
| ISIS EM-BLASSO | 2 | 8062916 | -2.70E-05 | 4.78 | 5.57 | 0.00 | 0.38 |
| pKWmEB | 2 | 8232320 | 1.00E-04 | 4.68 | 5.46 | 0.00 | 0.31 |
| ISIS EM-BLASSO | 2 | 8244606 | 3.00E-04 | 8.45 | 9.35 | 0.05 | 0.47 |
| pKWmEB | 2 | 8502325 | -5.00E-04 | 17.45 | 18.50 | 0.58 | 0.14 |
| ISIS EM-BLASSO | 3 | 72709 | 8.00E-04 | 7.14 | 8.01 | 0.14 | 0.14 |
| pKWmEB | 3 | 179032 | 1.00E-03 | 6.83 | 7.69 | 2.37 | 0.31 |
| ISIS EM-BLASSO | 3 | 179032 | 0.0025 | 24.04 | 25.16 | 2.60 | 0.30 |
| ISIS EM-BLASSO | 3 | 4095333 | 0.0013 | 11.05 | 12.01 | 0.74 | 0.33 |
| ISIS EM-BLASSO | 3 | 6640551 | -0.006 | 6.58 | 7.43 | 3.39 | 0.05 |
| mrMLM | 3 | 6640556 | -0.0103 | 5.46 | 6.28 | 10.10 | 0.05 |
| ISIS EM-BLASSO | 3 | 6640556 | -0.003 | 6.23 | 7.07 | 0.85 | 0.05 |
| pKWmEB | 3 | 6679615 | -4.00E-04 | 5.34 | 6.15 | 0.92 | 0.06 |
| ISIS EM-BLASSO | 3 | 6679615 | -6.00E-04 | 16.24 | 17.28 | 0.04 | 0.07 |
| pKWmEB | 3 | 7294460 | 4.66E-05 | 4.57 | 5.35 | 0.57 | 0.34 |
| ISIS EM-BLASSO | 3 | 7346278 | 6.00E-04 | 3.67 | 4.40 | 0.12 | 0.20 |
| pKWmEB | 3 | 7521430 | 3.00E-04 | 3.27 | 3.98 | 0.10 | 0.09 |
| ISIS EM-BLASSO | 4 | 25804 | -0.0019 | 10.68 | 11.63 | 1.31 | 0.25 |
| pKWmEB | 4 | 329152 | -1.30E-03 | 17.54 | 18.60 | 3.40 | 0.05 |
| pKWmEB | 4 | 2203719 | -1.00E-04 | 8.25 | 9.15 | 0.02 | 0.31 |
| pKWmEB | 4 | 3862627 | -3.00E-04 | 3.27 | 3.98 | 0.47 | 0.08 |
| ISIS EM-BLASSO | 4 | 4117472 | 4.00E-04 | 4.43 | 5.20 | 0.04 | 0.13 |
| pKWmEB | 4 | 4362316 | 1.00E-04 | 4.83 | 5.62 | 0.30 | 0.24 |
| ISIS EM-BLASSO | 4 | 4545038 | -0.0018 | 4.01 | 4.76 | 0.60 | 0.11 |
| pKWmEB | 4 | 5602779 | 6.59E-05 | 21.15 | 22.24 | 1.87 | 0.08 |
| ISIS EM-BLASSO | 4 | 7533605 | -2.00E-04 | 3.67 | 4.40 | 0.01 | 0.09 |

**Table 2.** List of QTNs significantly associated with tebuconazole.

| Method | Chromosome | Marker position (bp) | QTN effect | LOD score | '-log10(P)' | r2 (%) | MAF |
| --- | --- | --- | --- | --- | --- | --- | --- |
| pKWmEB | 1 | 3885008 | 9.54E-05 | 4.11 | 4.87 | 0.26 | 0.34 |
| ISIS EM-BLASSO | 1 | 5731674 | -5.00E-04 | 3.59 | 4.32 | 0.15 | 0.09 |
| pKWmEB | 1 | 5933539 | 9.64E-05 | 18.95 | 20.02 | 0.55 | 0.47 |
| ISIS EM-BLASSO | 1 | 7467655 | -0.001 | 11.23 | 12.20 | 0.39 | 0.05 |
| pKWmEB | 1 | 11258195 | 3.68E-05 | 24.88 | 26.01 | 2.08 | 0.09 |
| pKWmEB | 1 | 11270238 | 1.00E-04 | 17.78 | 18.84 | 0.19 | 0.22 |
| pKWmEB | 2 | 501355 | -7.00E-04 | 15.62 | 16.66 | 0.65 | 0.10 |
| ISIS EM-BLASSO | 2 | 665406 | 0.001 | 13.94 | 14.95 | 1.71 | 0.30 |
| ISIS EM-BLASSO | 2 | 851037 | 0.0014 | 12.02 | 12.99 | 2.83 | 0.24 |
| pKWmEB | 2 | 964956 | 5.68E-05 | 5.07 | 5.87 | 0.16 | 0.33 |
| pKWmEB | 2 | 985340 | 8.95E-05 | 11.02 | 11.98 | 2.06 | 0.17 |
| ISIS EM-BLASSO | 2 | 2645857 | -6.00E-04 | 3.94 | 4.69 | 0.46 | 0.22 |
| ISIS EM-BLASSO | 2 | 3938384 | 7.00E-04 | 3.20 | 3.91 | 0.80 | 0.33 |
| ISIS EM-BLASSO | 2 | 5182922 | 6.00E-04 | 3.31 | 4.02 | 0.26 | 0.10 |
| ISIS EM-BLASSO | 2 | 6143802 | 0.0011 | 3.79 | 4.53 | 1.23 | 0.15 |
| ISIS EM-BLASSO | 2 | 6173074 | 7.00E-04 | 8.11 | 9.01 | 0.50 | 0.18 |
| pKWmEB | 2 | 8129795 | 5.58E-05 | 3.20 | 3.91 | 0.87 | 0.35 |
| ISIS EM-BLASSO | 2 | 8222086 | -0.0013 | 7.99 | 8.88 | 2.26 | 0.22 |
| pKWmEB | 2 | 8322036 | -7.72E-05 | 30.42 | 31.59 | 3.06 | 0.35 |
| pKWmEB | 2 | 8566172 | -3.00E-04 | 13.45 | 14.45 | 1.13 | 0.06 |
| ISIS EM-BLASSO | 2 | 8566172 | -5.00E-04 | 6.71 | 7.56 | 0.11 | 0.06 |
| pKWmEB | 2 | 8880263 | -4.00E-04 | 3.47 | 4.19 | 0.00 | 0.18 |
| ISIS EM-BLASSO | 3 | 65770 | 4.00E-04 | 4.36 | 5.13 | 0.28 | 0.45 |
| pKWmEB | 3 | 503820 | 6.97E-05 | 12.29 | 13.28 | 0.12 | 0.34 |
| ISIS EM-BLASSO | 3 | 3183025 | 4.00E-04 | 4.07 | 4.82 | 0.25 | 0.33 |
| pKWmEB | 3 | 3813789 | 3.00E-04 | 32.56 | 33.75 | 0.57 | 0.07 |
| ISIS EM-BLASSO | 3 | 6299555 | 6.00E-04 | 5.12 | 5.92 | 0.19 | 0.09 |
| pKWmEB | 3 | 6562010 | 7.63E-05 | 16.65 | 17.69 | 1.61 | 0.15 |
| ISIS EM-BLASSO | 3 | 6640551 | -6.00E-04 | 3.00 | 3.70 | 0.12 | 0.05 |
| ISIS EM-BLASSO | 3 | 6679615 | -5.00E-04 | 6.09 | 6.93 | 0.12 | 0.07 |
| ISIS EM-BLASSO | 3 | 7066756 | -8.00E-04 | 12.90 | 13.89 | 1.02 | 0.35 |
| ISIS EM-BLASSO | 3 | 7067865 | -3.03E-05 | 4.73 | 5.52 | 0.00 | 0.43 |
| pKWmEB | 3 | 7346278 | 1.00E-03 | 32.95 | 34.14 | 2.53 | 0.21 |
| ISIS EM-BLASSO | 3 | 7346278 | 0.0012 | 14.90 | 15.92 | 1.76 | 0.20 |
| ISIS EM-BLASSO | 4 | 25804 | -0.0013 | 13.20 | 14.20 | 2.36 | 0.25 |
| ISIS EM-BLASSO | 4 | 177976 | -5.00E-04 | 6.82 | 7.68 | 0.12 | 0.08 |
| ISIS EM-BLASSO | 4 | 2203719 | -6.00E-04 | 5.20 | 6.00 | 0.54 | 0.31 |
| pKWmEB | 4 | 4545038 | -1.00E-03 | 10.08 | 11.02 | 1.23 | 0.11 |
| pKWmEB | 4 | 5201402 | -2.00E-04 | 12.27 | 13.25 | 0.31 | 0.05 |

## Discussion

Our use of a published high-throughput method for the measurements of EC_50_ for two DMI fungicides proved effective at detecting variation in our sample. We estimated a broad sense heritability of above 0.8 for sensitivity to each fungicide, while (Talas and McDonald, 2015) also reported high heritability of propiconazole sensitivity (0.97) in laboratory experiments conducted using essentially the same protocol. The distribution of our propiconazole EC_50_ values was roughly similar to those of (Talas and McDonald, 2015), who measured a range of 16 - 182 *μ*M and mean of 65 *μ*M. Our results are within a factor of two lower. In contrast, our measurements of EC_50_ for tebuconazole stand in greater contrast to those of (Spolti et al., 2014) and (Poudel et al., 2024). The sample of (Spolti et al., 2014) had a mean of 3.6 *μ*M, with a range of 0.9 - 26 *μ*M, while that of (Poudel et al., 2024) had a median of 2.8 *μ*M and range of 0.2 - 10 *μ*M. Our mean value is about five or six-fold larger than these previous studies. And this difference is not likely explained by differences in the sample alone, as our tebuconazole EC_50_ for the PH-1 isolate (11157), 11.75 *μ*M, is also about five times higher than Poudel *et al*.’s median measurement for the same isolate (Poudel et al., 2024). The reason for the discrepancy is unclear, and may be due to differences in experimental protocols or EC_50_ estimation. Nevertheless, our main focus was to obtain a series of precise, rather than accurate, measurements, since for the purpose of GWAS we only need relative values among isolates in our sample.

Our isolates were collected over a span of 14 years, with roughly half of the isolates collected in 2000 or earlier, and half collected in 2011 or 2013. While it is notable that the two isolates with the lowest sensitivity to both fungicides were collected in 2013, aside from these outliers, the EC_50_ distributions from the two time periods were similar for each fungicide. A more detailed analysis of shifts in fungicide sensitivity over time in the U.S. would require more isolates sampled from a broader time range, including isolates collected more recently than 2013.

Our GWAS scans identified a total of 48 significant QTNs for propiconazole sensitivity and 39 for tebuconazole sensitivity. Among the 72 distinct QTNs associated with azole sensitivity in this study, only 22 explain at least one percent of the phenotypic variation observed for either fungicide, and only a single QTN could explain at least 10% of the phenotypic variation. This indicates that, in the NA1 population of *F. graminearum,* azole sensitivity, as opposed to azole resistance, is governed by many QTNs with small effect. A couple of other studies have performed GWAS for fungicide sensitivity in *F. graminearum* populations. (Talas et al., 2016) reported 74 significant QTNs for propiconazole sensitivity in German field populations of *F. graminearum,* where each QTN explained up to 17% of the genetic variance in azole sensitivity. We identified three candidate genes (one for tebuconazole and two for propiconazole) common to the ones reported by (Talas et al., 2016). FGRAMPH1_01G13811 (FGSG_16193, FGSG _3830) is a quinate transporter, gene FGRAMPH1_01G19355 (FGSG _06044) is a transcription initiation factor TFIID subunit 12 and FGRAMPH1_01G22503 (FGSG _12935) is a hypothetical protein (Supplemental Table 2). In a similar study, (Poudel et al., 2024) reported one QTN associated with tebuconazole sensitivity that explained 48% of the phenotypic variation and six QTN associated with prothioconazole sensitivity, which explained from 2 to 16% of the phenotypic variation. The gene FGRAMPH1_01G15717, which was associated with prothioconazole sensitivity by (Poudel et al., 2024), was significantly associated with tebuconazole sensitivity in the current study. We expect in some cases that the same genetic variants will influence sensitivity to multiple azole fungicides because of their shared mode of action, but because different azole fungicides show differential binding preference to different CYP51 paralogs in *F. graminearum*, we also expect GWAS scans may detect variants unique to a single azole fungicide (Liu et al., 2011). Additionally, outside of the CYP51 paralogs, other variants may have different effects on a strain’s sensitivity to distinct fungicides. The single candidate gene for tebuconazole sensitivity observed by (Poudel et al., 2024) was not identified in this study. This could be due to differences between the studies in the composition of the samples of isolates and their different geographic origins. While the Poudel *et al*. study included only isolates collected in North Dakota, the population structure apparent from that study and presence of a large proportion of 3ADON isolates indicates that their samples likely represent isolates from the NA1 and NA2 populations, if not others. And the NA1 and NA2 populations have fairly distinct evolutionary histories, with many variants found at greatly different frequencies between the populations (Dhakal et al., 2024). Thus, SNPs found associated by Poudel *et al*. could represent differences in fungicide sensitivity between populations. In comparison, over 96% of the isolates included in this study are confidently assigned to the NA1 population, such that the QTNs we identify must be variable within this population. Despite the differences in the samples used in each study, candidate genes common between our study and those of (Talas et al., 2016) and (Poudel et al., 2024) provide supporting evidence for the involvement of these genes, which represent strong candidates for functional characterization. Similar to (Talas et al., 2016), we did not identify any genes overlapping with those significantly differentially expressed in response to tebuconazole treatment reported by (Becher et al., 2011).

One of our candidate genes, FGRAMPH1_01G09621 (FGSG _17058), is a multidrug resistance type ABC transporter and has been previously identified to specifically play a role in tolerance to SBI class I triazoles (tebuconazole, triticonazole and epoxiconazole) but not to the SBI type II triazoles (Abou Ammar et al., 2013; Goswami et al., 2006). The gene was identified as a candidate gene in tebuconazole but not propiconazole sensitivity in our analyses. Our results further support that different ABC transporters might be involved in tolerance to different fungicides. In addition to this multidrug resistance protein, we identified five other transport related proteins (FGRAMPH1_01G13811, FGRAMPH1_01G16125, FGRAMPH1_01G19393, FGRAMPH1_01G22013, and FGRAMPH1_01G25701) among our candidate genes. Transport of fungicide out of the cells is known to be one of the molecular mechanisms for tolerance or resistance to fungicides in fungi (Hayashi et al., 2003; Reimann and Deising, 2005), and our results indicate that *F. graminearum* may also have this mechanism in place for fungicide tolerance.

Interestingly, genes within the fusarin C biosynthetic cluster were identified among the candidate loci associated with sensitivity to propiconazole and tebuconazole. Sublethal fungicide doses are known to stimulate secondary metabolite production (Felix D’Mello et al., 1998) and we speculate that the transporter gene within the fusarin C cluster may contribute to enhanced tolerance to these DMIs. Supporting this hypothesis, *Penicillium expansum* isolates resistant to multiple fungicides have been shown to produce higher levels of patulin and exhibit increased expression of efflux transporters linked to patulin biosynthesis (Ntasiou et al., 2023). Similarly, *Botrytis cinerea* infection in grapes induces plant secondary metabolite production and overexpression of ABC transporters, which in turn drives multidrug resistance in the pathogen (Hayashi et al., 2003). Beyond the fusarin C cluster, other secondary metabolite biosynthetic genes, including nonribosomal peptide synthetases (FGRAMPH1_01G08375, FGRAMPH1_01G20955) and a polyketide synthase (FGRAMPH1_01G08383) were also identified as candidate genes potentially involved in azole sensitivity. We hypothesize that *Fusarium graminearum* employs transporters both within secondary metabolite gene clusters and outside these clusters to actively remove fungicides from the cell.

Heat shock proteins (HSPs) help fungi respond to heat and other types of environmental stressors. HSP90 and HSP70 are also responsible for antifungal resistance in *Candida albicans* and other fungi (Blatzer et al., 2015; Cowen and Lindquist, 2005; Lamoth et al., 2015). HSP70 (FGRAMPH1_01G21231) is a novel candidate gene identified for both propiconazole and tebuconazole sensitivity in this study. The role of HSP70 in azole resistance should be investigated further.

We did not identify any obviously resistant isolates in our study. Mutations in CYP51 genes have been related to azole resistance in many fungi, including *Mycosphaerella graminicola* (Leroux et al., 2007). However, we did not identify any significant associations with variants in or near CYP51 genes in *F. graminearum*. This could be due to the following reasons: i) field isolates of *F. graminearum* from our sample may not have mutations in CYP51 genes that lead to resistance, and ii) genes outside of CYP51 loci may contribute the largest portion to variation in fungicide tolerance. (Talas and McDonald, 2015) considered the latter as one of the explanations for no observed correlation between mutations in CYP51 genes and propiconazole resistance in *F. graminearum*. Our results show that genes outside CYP51 loci are responsible for variation in azole tolerance in our sample, although functional validation remains to be done. In our analysis, we identified seven transcription factors as candidate genes for propiconazole and tebuconazole sensitivity. Overexpression of CYP51 genes is another known mechanism for tebuconazole resistance in fungi (Hamamoto et al., 2000; Ma et al., 2006). We can speculate that overexpression of other genes like the transporters can lead to fungicide tolerance or resistance, and these transcription factors could be affecting the expression levels of genes important in azole sensitivity.

We did not detect any *Fusarium graminearum* isolates that we would classify as resistant to either propiconazole or tebuconazole. However, due to the high genetic diversity and frequent sexual reproduction in *F. graminearum* populations, there remains significant potential for the development of fungicide resistance. Therefore, continued monitoring of populations is essential to detect shifts in fungicide sensitivity over time. The candidate genes identified in this study provide valuable targets for future functional characterization and may help elucidate the molecular mechanisms underlying DMI tolerance.

## Acknowledgments

We are grateful to Gary Bergstrom, H. Corby Kistler, John F. Leslie, and Shaobin Zhong for providing the isolates used in this study. We thank Paul St. Amand for his assistance with the Biomek liquid handling automated workstation. This is contribution no. 26-030-J from the Kansas Agricultural Experiment Station.

## Author contributions

U.D.: Methodology, Investigation, Formal analysis (lead), Data curation, Validation, Visualization, Writing - original draft, Writing - review & editing

C.T. Conceptualization, Funding acquisition, Project administration, Formal analysis (secondary), Supervision Writing - review & editing

---

**Supplemental Figure 1:**
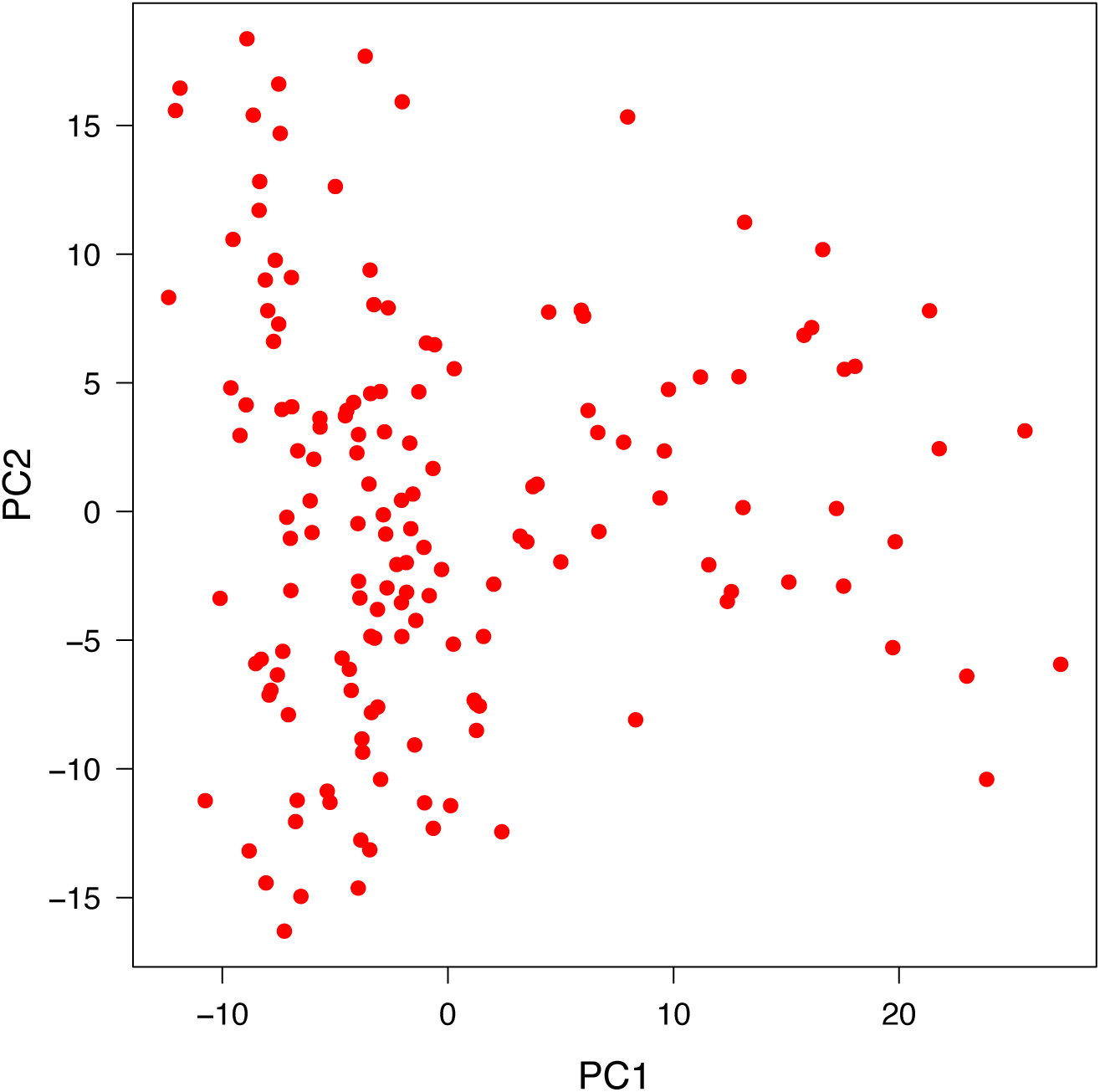
Principal component analysis of 152 *F. graminearum* isolates used in this study. Isolates are plotted along the first (PC1) and second (PC2) principal components of SNP variation.

